# Reproducible and sensitive micro-tissue RNA-sequencing from formalin-fixed paraffin-embedded tissue for spatial gene expression analysis

**DOI:** 10.1101/2022.03.29.486169

**Authors:** Hiroko Matsunaga, Koji Arikawa, Miki Yamazaki, Ryota Wagatsuma, Keigo Ide, Samuel Ashok Zachariah, Kazuya Takamochi, Kenji Suzuki, Takuo Hayashi, Masahito Hosokawa, Hideki Kambara, Haruko Takeyama

## Abstract

Spatial transcriptome analysis of formalin-fixed paraffin-embedded (FFPE) tissues using RNA-sequencing (RNA-seq) provides interactive information on morphology and gene expression, which is useful for clinical applications. However, despite the advantages of long-term storage at room temperature, FFPE tissues may be severely damaged by methylene crosslinking and provide less gene information than fresh-frozen tissues. In this study, we proposed a sensitive FFPE micro-tissue RNA-seq method that combines the punching of tissue sections (diameter: 100 μm) and the direct construction of RNA-seq libraries. We evaluated a method using mouse liver tissues at 2 years after fixation and embedding and detected approximately 7,000 genes in micro-punched tissue-spots (thickness: 10 μm), similar to that detected with purified total RNA (2.5 ng) equivalent to the several dozen cells in the spot. We applied this method to clinical FFPE specimens of lung cancer that had been fixed and embedded 6 years prior, and found that it was possible to determine characteristic gene expression in the microenvironment containing tumor and non-tumor cells of different morphologies. This result indicates that spatial gene expression analysis of the tumor microenvironment is feasible using FFPE tissue sections stored for extensive periods in medical facilities.

## Introduction

Integrated analysis of cell spatial location and gene expression information using histological tissue sections is a powerful tool for determining tissue function, particularly pathological changes, under diseased conditions^1–7^. Initially, hybridization-based methods, such as FISH or MERSFISH, were developed for the spatial analysis of genes expressed *in vivo* using fluorescent probes prepared in advance^1–3^. However, issues such as limited adaptation to only known genes and limited simultaneous detection of multiple genes were reported. With the rapid progress of next-generation sequencing technology, spatial transcriptome technology has recently garnered attention for comprehensive sequence-based analysis^4,6,7^. All tissue functions involve complex cell–cell interactions^8–10^. Examining the gene expression profiles of morphologically specific tissue domains could help determine the details of cellular interactions in such microenvironments. Fresh-frozen (FF) tissue-based sampling is widely used for mainstream gene expression analyses^1–4,6,7^, owing to the relative ease of extraction of high-quality RNA. However, it is difficult to preserve FF tissue samples for long periods, and the method does not allow detailed morphological observation.

Morphological characteristic-based tissue diagnostic approaches are valuable in the clinical field^11–13^. Cellular structures and tissue morphologies are generally well-preserved in formalin-fixed paraffin-embedded (FFPE) tissues, and hence, they are preferred over FF tissues for sample preservation in pathological diagnosis. In addition, the potential for long-term storage at room temperature is an important advantage of FFPE samples. Archived FFPE tissue samples often serve as invaluable resources for diagnosing disease pathogenesis. However, the crosslinking of proteins and nucleic acids may occur during formalin fixation and cause RNA fragmentation^14,15^. This in turn affects the quality of FFPE-derived RNA; hence, whole-transcriptome analysis of FFPE tissues using standard protocols is challenging. Recent studies have examined the feasibility of transcriptome analysis using purified total RNA from FFPE tissues and also compared the gene expression in FF and FFPE tissues^16–20^. However, it was challenging to obtain gene expression information using methylene-crosslinked RNA derived from FFPE tissues, and each method required a pre-treatment process involving extraction and purification from tissue sections of a specific size (at least 5 mm^2^ approximately) used as the starting material with a commercially available kit.

Spatial transcription profiling is a powerful tool for evaluating the functional characteristics of cells in a typical tissue architecture. It is required to focus on minute, specific regions for understanding the interaction between tissue morphology and corresponding gene expression. Obtaining gene expression information with minimum loss of RNA molecules from a very small tissue section is critical for the success of spatial transcriptomic analysis. In a previous study, we targeted very small (< 100 μm) micro-tissues containing a few tens of cells for constructing RNA-sequencing (RNA-seq) libraries^21,22^. Typical kit-based RNA extraction protocols are not suitable for such small samples, as a certain quantity of RNA molecules are potentially lost during nucleic acid purification, which may involve the use of columns. It is desirable to develop a protocol that is compatible for use with small tissue sections, minimizes sample loss, and allows processes from RNA extraction to cDNA construction to be performed in an uncomplicated manner. Previously, we had reported the development of a high-speed apparatus for the punching and collection of micro-<u>pu</u>nched <u>ti</u>ssue <u>spots</u> (PuTi-spots) (diameter: 100 μm) from FF tissue slices (thickness: 10–20 μm) while observing the morphological characteristics under a microscope. Using this punching device, PuTi-spots could be collected from both FF and FFPE tissue sections.

In this study, we report gene expression analysis using very small FFPE tissue sections, a process that has not been conducted with much success to date. We developed an upgraded RNA-seq library preparation protocol for FFPE PuTi-spots using a surfactant, proteinase K, and heating for membrane dissolution and decrosslinking, based on a previously reported method^22^. We critically compared the detection sensitivity and accuracy of RNA-seq achieved using this method with those achieved using serially diluted purified total RNA as the starting material. Further, we performed a thorough comparison of the RNA-seq results obtained using FFPE and FF tissues. Finally, RNA-seq analysis of the PuTi-spots was performed using FFPE clinical specimens to evaluate the detection of genes in low-quality tissue samples with degraded of nucleic acids after long-term archiving at room temperature for more than 6 years.

## Results

### Sample summary

An overview of the samples used for the assessments in this study is provided in Figure 1a. Frozen mouse liver tissue stored at -80 °C for 2 years after embedding was used as the FF sample. Similarly, mouse liver tissue fixed in 4% formalin for 24 h, embedded in paraffin, and stored at 4 °C for 2 years was used as the FFPE sample. In both cases, purified total RNA was analyzed using thin tissue sections (thickness: 10 μm) with commercially available RNA extraction kits. Total RNA quality was assessed based on the RNA integrity number (RIN)^23,24^ estimated by electropherograms using Tape Station (Agilent Technologies, Santa Clara, CA, USA) (Table 1). For the FF sample, 18S and 28S rRNA peaks could be clearly identified, and the average RIN value was estimated to be 8.4 (N = 3) (Supplementary Figure 1). Conversely, the FFPE sample showed no observable rRNA peaks, and the average RIN value was 3.5 (N = 3), which was significantly lower than that of the FF sample. To test the possibility of RNA degradation, the DV200 values (percentage of RNA fragments with > 200 nucleotides) were calculated based on the results of electropherograms^25^. The DV200 value for the FF sample was 98.3%, indicating almost no degradation. The DV200 value for the FFPE sample was 86.3%, which was relatively lower.

**Table 1.** Sample quality assessments.

|  | Preparation | N | RNA quality |  |
| --- | --- | --- | --- | --- |
|  |  |  | RIN | Dv200 |
| Fresh frozen | 2017_09 | 3 | 8.4 | 98.3% |
| FFPE | 2017_07 | 3 | 3.5 | 86.3% |

RIN: RNA integrity number
DV200: Percentage of RNA fragments >200 nucleotides

**Figure 1.**
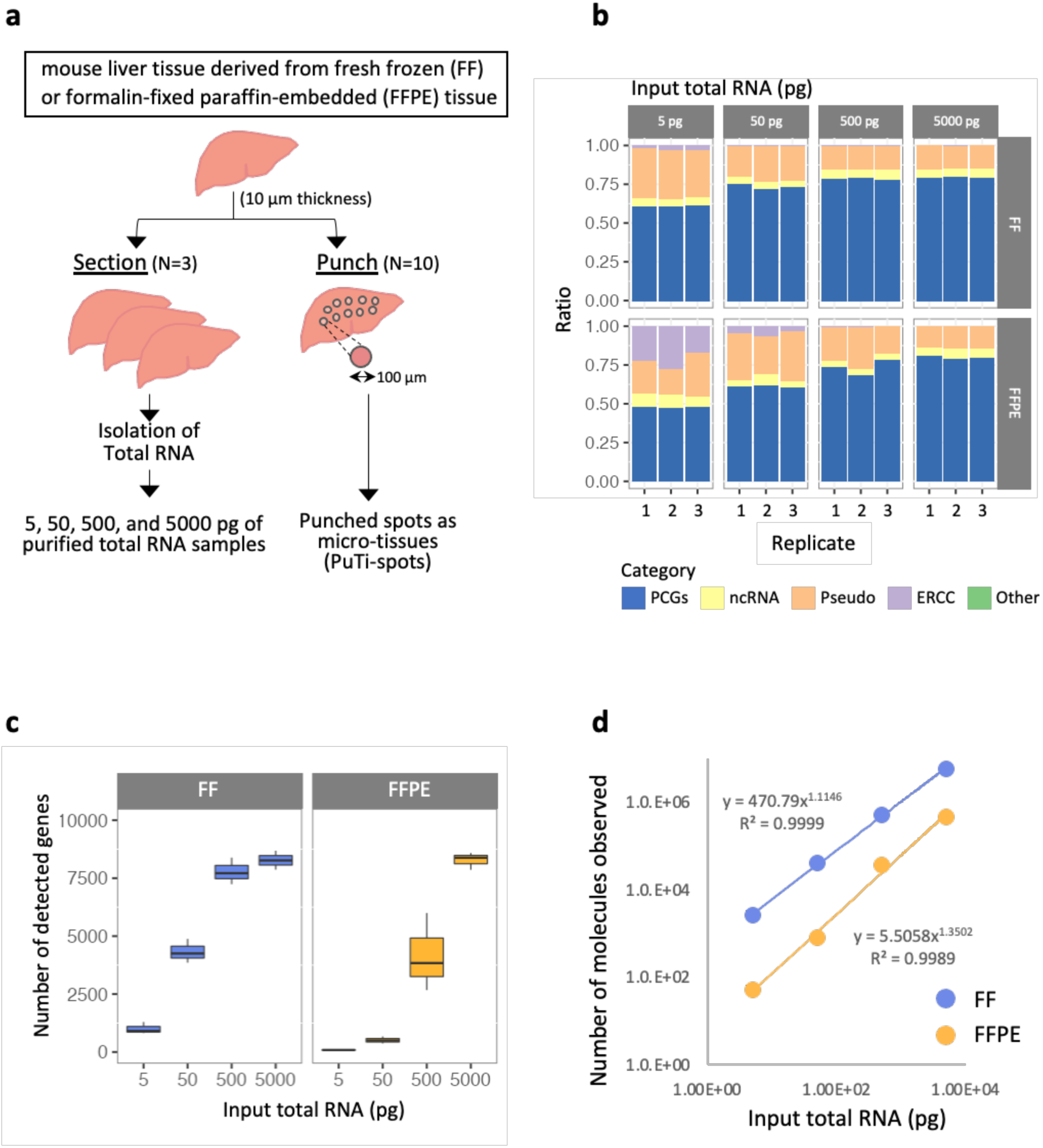
Evaluation of RNA-seq profiling of purified total RNA derived from FFPE and FF tissues. (a) Overview of sample preparation for evaluation using mouse liver tissue. Purified total RNA samples were prepared using the RNeasy Mini Kit (QIAGEN) for FF tissues or RNeasy FFPE Kit (QIAGEN) for FFPE tissues. (b) Mapping category of reads. We evaluated whether reads were assigned to five categories: PCGs, ncRNAs, pseudo genes, ERCC of internal standard RNAs, or others. As the input increased, the ratio assigned to the PCGs increased, and above a certain quantity, the values were saturated. It is also apparent that the assignment of reads to ERCC increases when PCGs have fewer reads, starting at 5 or 50 pg. (c) Number of detected genes. The number of genes detected for respective input RNAs was plotted. Libraries were prepared using total RNA purified from FF or FFPE tissue sections (N = 3) as the starting material. PCGs were identified from the respective libraries. (d) Number of RNA molecules from total RNA contributing to the reaction. We estimated the number of RNA molecules that contribute to the reaction with reference to ERCC, which was used as an internal standard, that is, the value of (sum of TPMs of the RNA) × (sum of the number of molecules of the input ERCC)/(sum of TPMs of the ERCC) = (sum of the number of input RNA molecules) was plotted. FFPE, formalin-fixed paraffin-embedded; FF, fresh frozen; RNA-seq, RNA-sequencing; PCGs, protein-coding genes; ncRNAs, non-coding RNAs; TPM, transcripts per kilobase million.

PuTi-spots were collected using the punching device^21^ from similar sliced thin sections (of FF or FFPE tissues). We collected 10 PuTi-spot (φ = 100 μm) replicas from one tissue section. Although it is considered that there are only few changes in gene expression at different sites in the liver^26^, adjacent tissue slices were used as replicas to minimize the variations in gene expression between samples owing to spatial heterogeneity (N = 3). For the same reason, adjacent spots collected from the same section with the punching device^21^ were used as PuTi-spot replicas (N = 10).

### Evaluation of RNA-seq profiles using purified total RNA from FF and FFPE tissue specimens

First, we performed RNA-seq of FFPE and FF tissue samples, beginning with different initial purified total RNA contents, to assess the potential impact of RNA degradation on the results of gene expression analysis. Four purified total RNA samples with 5, 50, 500, and 5,000 pg (= 5 ng) of total input RNA per reaction were prepared by serial dilution. The smallest input total RNA content of 5 pg per reaction was selected because the total RNA content per cell is expected to range from 5 to 50 pg. PuTi spots are assumed to contain 10-50 cells based on the diameter and thickness of the spot and the nuclear staining pattern. Therefore, a series of total RNA reaction mixtures with RNA content of up to 5,000 pg was used to match the RNA content in the PuTi-spot. Three replicates were used for each case (N = 3). The average number of sequence-reads per sample was 0.6 million. The proportion of reads assigned to protein-coding genes (PCGs) in FF samples plateaued at 80% of the total reads (Figure 1b). In FFPE, the ratio of PCGs reached 80% at 5,000 pg of input. Compared to that in FF samples, the value in FFPE samples increased to 80%, with an order of magnitude higher amount of inputs (5,000 pg). The number of genes detected in the FF sample also plateaued at an input of 500 pg, whereas the genes detected in the FFPE sample reached a similar number at an input of 5,000 pg, which was at a higher order of magnitude (Figure 1c). This indicates that between FF- and FFPE-derived samples, owing to RNA degradation, the actual number of RNA molecules in the mixture participating in the reaction could be different from the total RNA input.

We then estimated the number of RNA molecules that contributed to cDNA production. The calculations were performed using ERCC RNA Spike-Ins (Thermo Fisher Scientific, Waltham, MA, USA), which is a commercially available external RNA control, and calculated values of transcripts per kilobase million (TPM). According to our estimate, we assumed that the ratio of the total number of ERCC molecules used in the reaction to the total TPM of ERCC (obtained from the RNA-seq results) is equal to the ratio of the total number of RNA molecules contributing to cDNA production to the total TPM of RNA. The number of molecules presumed to have contributed to the reaction increased by an order of magnitude relative to the abundance ratio of the dilution series of total RNA from 5 pg to 5,000 pg used for the reaction (Figure 1d). This was anticipated, implying that the number of molecules contributing to the reaction increases with the total RNA used in the reaction. Conversely, the estimated number of molecules contributing to the reaction in FF and FFPE tissues differed by almost an order of magnitude (Figure 1d). This implies that the reads assigned to PCGs plateaued at a value less than that in the FF samples by an order of magnitude because the number of RNA molecules contributing to the reaction in the FF-derived purified total RNA was greater by one order of magnitude. This result suggests that the 3’-end polyA sequence of the mRNA was not sufficiently captured by the poly-T primer used in the reverse transcription reaction owing to progressive degradation or incomplete decrosslinking of the purified total RNA molecules from FFPE tissues. The FFPE tissue samples used in this study were formalin-fixed for 24 h and then paraffin-embedded and stored at 4 °C for 2 years. The dissociation of methylene crosslinking was challenging, even for tissues that had been stored under relatively mild conditions (4 °C). In the commercial kit used in this study, decrosslinking was performed by heat treatment. Although heat treatment is known to cause the effective dissociation of methylene crosslinking, it may also induce RNA degradation. Even when the total RNA derived from FFPE tissue was purified by the optimization protocol recommended in the kit, the reaction efficiency was lower than that achieved with the total RNA derived from FF tissue, corresponding to a reduction by an order of magnitude in the input volume.

### RNA-seq analysis of PuTi-spots from mouse liver specimens

RNA-seq analyses of mouse liver PuTi-spots were performed for validating the sequence quality of the FFPE samples using our proposed method. The micro-punched circular sections were estimated to contain 10 to 50 cells per spot, as determined by the counting of hematoxylin-eosin (HE)-stained nuclei. An overview of the workflow from PuTi-spot collection to RNA-seq library preparation is shown in Figure 2a. The FFPE tissue sections used in this study were fixed in formalin for 24 h, paraffin-embedded, and stored at 4 °C for 2 years, as described above. After deparaffinization of the FFPE tissue slices, the regions of interest in the tissue were selected based on morphology and micro-punched using our sampling device^21,22^. FF tissue was also used for evaluation after 2 years of storage at -80 °C embedded in super cryoembedding medium (SCEM) solution (Leica Microsystems, Tokyo, Japan). Before micro-punching, the FF tissue sections were lightly fixed in 99.5% ethanol immediately after sectioning to inhibit RNA degradation^22^. PuTi-spots were recovered directly into microtubes from both FFPE and FF samples and were then subjected to cell membrane disruption by proteinase K treatment and mRNA purification using polyT magnetic beads (PPP, <u>p</u>urification by <u>p</u>roteinase K and <u>p</u>olyT magnetic beads)^22^. For FFPE tissues, heat treatment was performed prior to magnetic bead treatment to induce the dissociation of methylene crosslinking. As previously reported, we confirmed that the PPP process is highly effective at improving the detection of PCGs in the cDNA synthesis of FF PuTi-spots^22^. In FFPE PuTi-spots, the number of genes detected without PPP treatment was 2,055 ± 269 (N = 4), whereas the number of genes detected with PPP involving permeabilization was 5,335 ± 325 (N = 4) (Figure 2b). The number of detected genes increased to 7,066±733 (N = 10) in response to heat treatment for the dissociation of methylene crosslinking.

**Figure 2.**
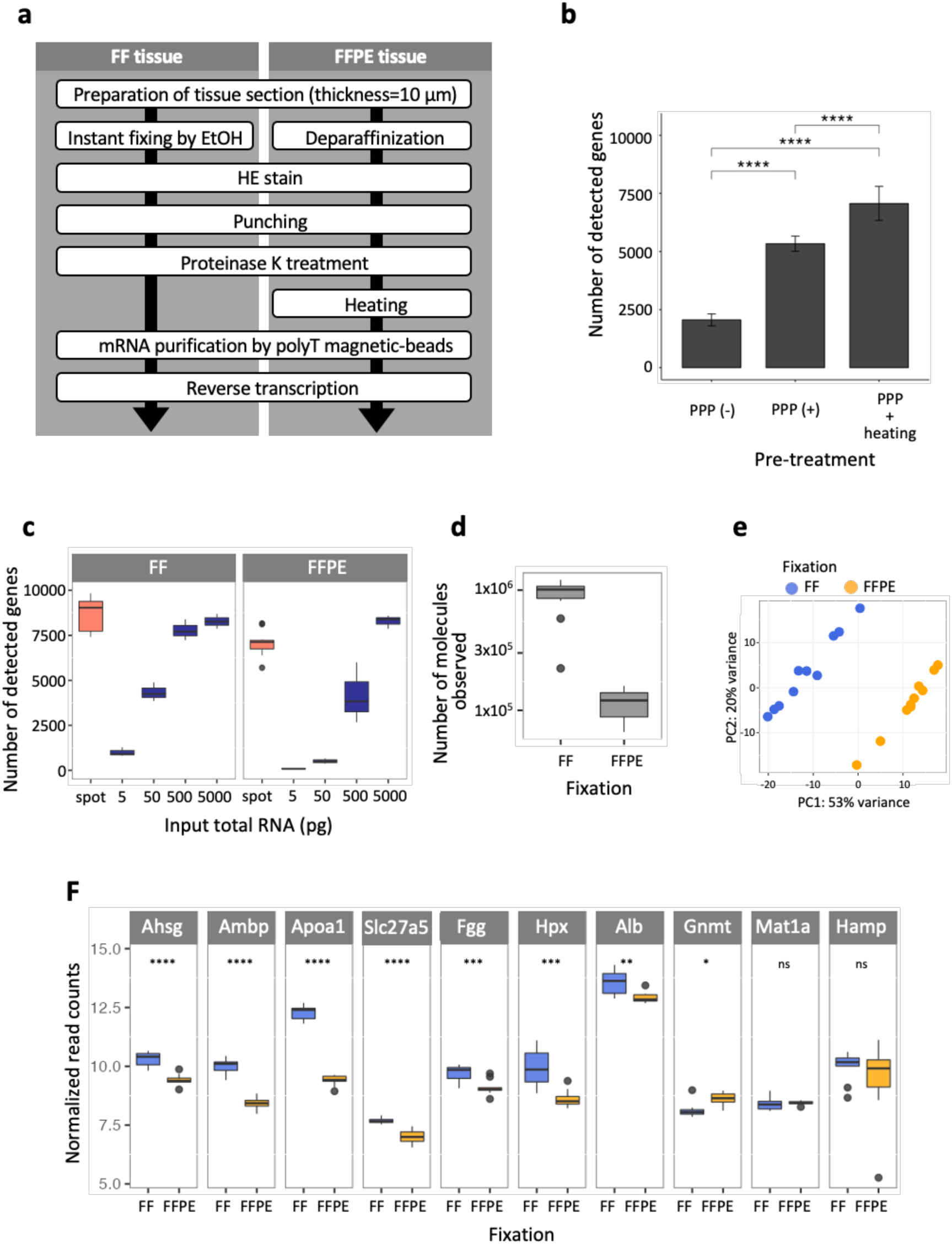
Evaluation of RNA-seq profiling data derived from FFPE and FF PuTi-spots. (a) Sample preparation flow of cDNA library derived from FF or FFPE PuTi-spots as starting materials. The difference between the handling of FF and FFPE tissues is related to the appropriate pre-treatment process for each fixation method before the punching of a micro-region and the additional heat treatment step for decrosslinking in the FFPE tissue. (b) Pre-treatment assessment for dissociation of crosslinking in FFPE PuTi-spots. (c) Comparison of the number of detected genes. The quantity of total RNA was assessed, which was equivalent to the number of genes detected in the micro-region samples. The number of genes detected from the micro-regions roughly corresponded to 500–5000 pg of total RNA. (d) As with the purified total RNA samples (Figure 1d), we estimated the quantity of RNA contributing to the reaction based on the molecular weight of ERCC used as an internal standard. (e) PCA of gene expression data. PCA was performed using gene expression data from FF or FFPE-derived PuTi-spots. Two different clusters were identified by PCA. The variance in percentages accounted for by each principal component is shown on each axis. (f) Expression levels of liver-specific genes observed using FF and FFPE PuTi-spots. We focused on ten genes specifically expressed in the mouse liver and compared the normalized read counts between FF and FFPE PuTi-spots used as starting materials (N = 10, respectively). Statistical analysis based on Welch’s *t*-test. *P value* < 0.05 (*), < 0.01 (**), < 0.001 (***), < 0.0001 (****). *P value* > 0.05 is indicated by “ns”. FFPE, formalin-fixed paraffin-embedded; FF, fresh frozen; PuTi-spot, micro-punched tissue spot; RNA-seq, RNA-sequencing; PCA, principal component analysis.

The average number of genes detected in one PuTi-spot was 8,688 for FF and 7,066 for FFPE samples. This corresponded to an input of 500 to 5,000 pg of total RNA in purified total RNA analysis (Figure 2c). Since the total RNA content per cell is approximately 50 pg, 50 cells would be roughly equivalent to a starting material of 2,500 pg of total RNA. The number of molecules contributing to the reaction was also calculated as previously performed for the purified total RNA sample. The estimated number of molecules contributing to the reaction was 10^6^ for FF and 10^5^ for FFPE samples, equivalent to a total RNA input of 2,500 pg (Figure 2d). In summary, we confirmed that using the proposed method, RNA present in the PuTi-spot can be recovered for RNA-seq analysis with efficiency comparable to that achieved with purified total RNA extraction using a commercially available kit.

### Evaluation of gene detection sensitivity in PuTi-spots of mouse liver specimens

The gene expression profile of the PuTi-spots was evaluated. Ten PuTi-spots were collected from a 10 µm-thick tissue section of mouse liver and used for evaluation as 10 replicates. The proportion of gene expression levels and detection frequency for both FF and FFPE samples were comparable in all 10 replicates (Supplementary Figure 2). In the principal component analysis (PCA) of gene expression, two different clusters corresponding to FF and FFPE samples were observed (Figure 2e). This suggests that the differences in the detected gene pattern depend on the tissue preservation condition. We compared the expression levels of 10 genes expressed specifically in the mouse liver^27^ in FF or FFPE tissue spots. The normalized read counts showed that the *Gnmt, Mat1a*, and *Hamp* genes showed almost no difference in expression between FF and FFPE samples, whereas the seven other genes showed significant differences in expression based on the fixation conditions (Figure 2f). The differences in the normalized read counts for each gene between PuTi-spots and purified total RNA were not substantial (Supplementary Figure 3), suggesting that the differences indicated in Figure 2f did not result from differences in the methods used for cDNA library preparation, including RNA purification. Therefore, we aligned the sequencing reads to each gene using Integrative Genomics Viewer^28–30^. The results showed that FFPE samples used for this evaluation showed a significant decrease in reads with alignment at approximately 500 bases from the 3’-end of each gene, regardless of whether the starting material was purified total RNA or a PuTi-spot. This suggests that cDNA obtained from FFPE samples is fragmented to approximately 500 bases or less owing to nucleic acid degradation by methylene crosslinking. Thus, the number of reads in FFPE-derived samples tends to decrease compared to that in the FF-derived sample because the reads are biased toward the 3’-end. Since the *Hamp* gene was only of approximately 400 bases, the reads mapped to the entire gene region without bias, which resulted in no difference between the normalized read counts in FF and FFPE samples. In contrast, the *Mat1a* gene spans approximately 3.5 kb, which is greater than the average length of the genes assessed in this study (approximately 1.6 kb). In both FF and FFPE samples, there was a depression in read alignment in the middle of the last exon at the 3’-end (the last exon was approximately 1.9 kb on length), possibly because the complete sequence was not obtained, similar to that obtained for the cDNA template, even from FF samples. The results of RNA-seq using PuTi-spots suggested that the differences in gene expression profiling could be attributed to the method of tissue preservation rather than the method of library preparation, based on the starting materials. The bias in gene expression profiling between FF- and FFPE-derived samples could be eliminated by, for example, focusing only on the 3′-end.

### RNA-seq analysis of PuTi-spots from tumor and non-tumor areas in resected pathological specimens of lung cancer

We applied the methods discussed above to a pathological specimen to evaluate their potential for clinical application. FFPE tissues derived from human lung cancer were used as the specimen, which had been embedded 6 years ago and were stored at room temperature. As in the previous analyses, total RNA was extracted from the thin sections (10 μm-thick) using a commercially available kit and used for RNA quality evaluation. In addition, PuTi-spots were prepared from sections adjacent to those used for RNA assessment. Punching was performed in each area, distinguishing between tumor and non-tumor areas, based on the discretion of the pathologist (Figure 3a). The RIN value was 1.8 and the DV200 value was 52%, which were indicative of progressive nucleic acid degradation by crosslinking.

**Figure 3.**
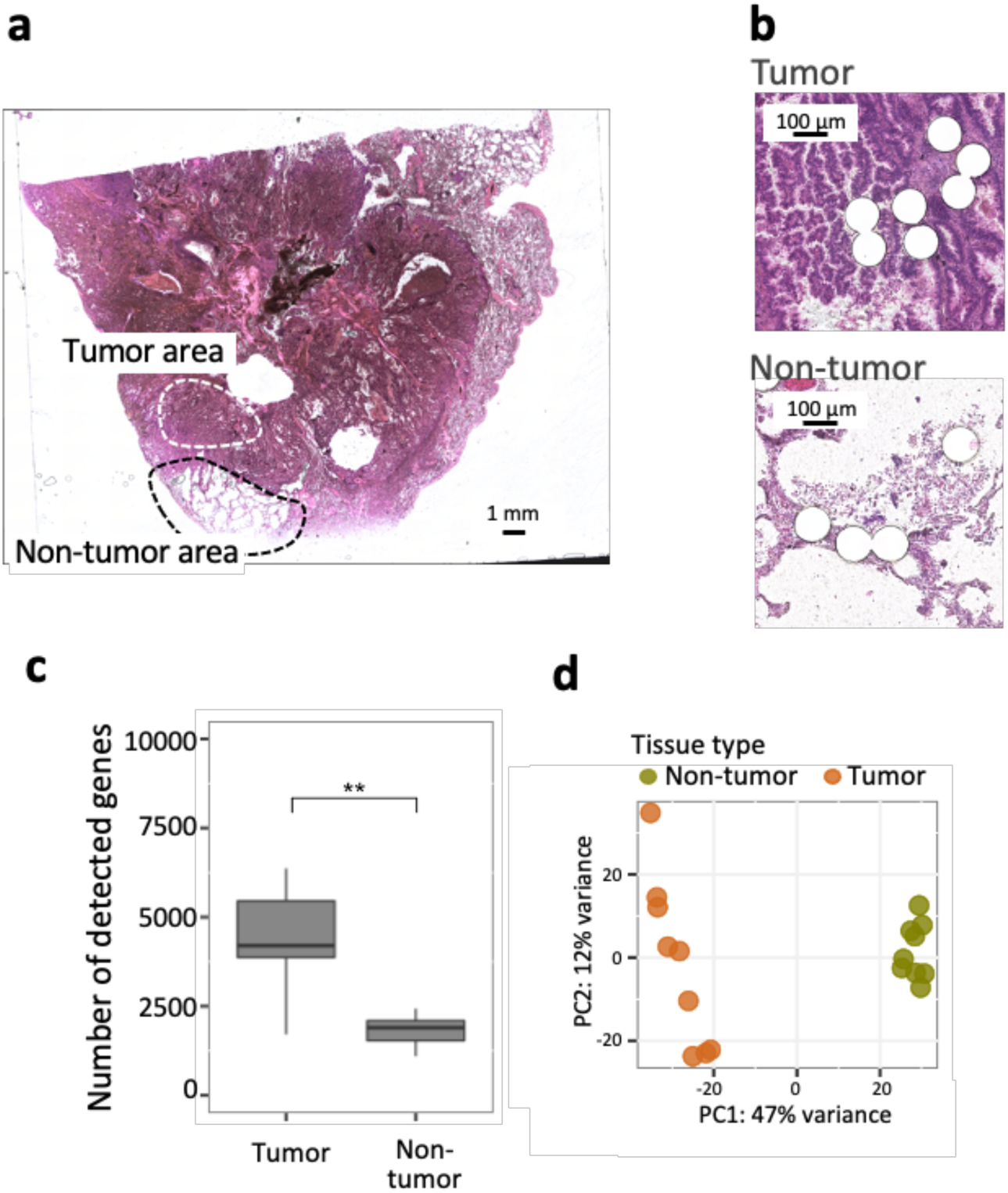
Evaluation of RNA-seq profiling of microenvironments in lung tumor specimens using PuTi-spots. (a) Morphological image of tumor and non-tumor areas in lung tumor specimens. Thin sliced sections (thickness: 10 μm) were obtained from a surgical resected sample that had been archived at room temperature for 6 years. After deparaffinization and HE staining, PuTi-spots were punched and collected under microscopic observation. (b) Image after PuTi-spot collection. Each white circle (diameter: 100 μm) corresponds to the location at which the PuTi-spot was punched out. (c) Comparison of the detected gene numbers. The quantity of total RNA samples was assessed, which was equivalent to the number of genes detected in the micro-region samples. The number of genes detected from the micro-regions roughly corresponded to 500–5000 pg of total RNA. Statistical analysis based on Welch’s *t*-test. \*\**P value* < 0.01 (d) PCA of gene expression data. PCA was performed using gene expression data from tumor or non-tumor-derived PuTi-spots (N = 10, respectively). Two different clusters were identified, and the distribution of the plots showed different trends between the groups. The amount of variance in percentages accounted for by each principal component is shown on each axis. RNA-seq, RNA-sequencing; PuTi-spot, micro-punched tissue spot; HE, hematoxylin-eosin; PCA, principal component analysis.

RNA-seq was performed using the PuTi-spots collected from the tumor and non-tumor areas (each N = 9) of the specimen (Figure 3b). The number of detected genes was greater in the tumor area than in the non-tumor area, with an average of 4,400 genes in the tumor area and 1,800 genes in the non-tumor area (Figure 3c). PCA showed that the gene expression pattern could be clearly categorized into two groups, one specific to the tumor area and the other to the non-tumor area (Figure 3d). Conversely, the variation among PuTi-spots was higher in the tumor area than in the non-tumor area. In this specimen, cadherin binding was extracted by enriched gene ontology (GO) analysis from among the genes upregulated in the tumor area (Supplementary Figure 4). Among the genes associated with cadherin binding, *S100P* and *TAGLN2* were found to exhibit upregulation (Supplementary Table 1). *S100P* is a Ca^2+^-binding protein that is overexpressed in various cancers and considered a tumor biomarker^31^. *TAGLN2* is a gene that has been suggested to be associated with cancer development, and suppressing this gene is considered to inhibit the growth, invasion, and metastasis of cancer cell^32^. Based on these results, it was indicated that even when PuTi-spots containing only a few dozen cells were collected from low-quality tissues with a DV200 value of 50%, a sufficient number of genes could be detected to understand the characteristics of cells with different morphologies, such as tumor and non-tumor cells, and to capture the different expression patterns functioning in the microenvironment.

## Discussion

We have demonstrated that direct RNA-seq from PuTi-spots (tissue spots with a diameter of 100 μm) derived from FFPE tissue stored for over 2 years can be used for gene expression analysis, with sensitivity comparable to that achieved with total RNA extracted using a commercially available kit. The key points of the featured experimental protocol are as follows. We demonstrated efficient membrane lysis and methylation crosslink dissociation for RNA extraction from the cells. For the FF tissue spots, membrane lysis was achieved using a surfactant and proteinase K, as reported previously^22^. For the FFPE tissue spots, we strengthened the process by using a surfactant, proteinase K, and heat treatment. Treatment with proteinase and heat was essential for the removal of proteins crosslinked with nucleic acids in the FFPE tissue. The library preparation protocol presented in this study can help minimize RNA loss by eliminating the need for RNA purification by column extraction. Consequently, we demonstrated that gene expression analysis can be performed with equal accuracy and sensitivity using mouse liver samples and purified total RNA, even in very small tissue sections with a diameter of 100 μm, which were collected from specific histomorphological locations. In previous analyses, the starting material was a thin section, with an area of approximately 5 mm^2^ or greater, cut from an FFPE tissue block, from which total RNA (in the order of micrograms) was extracted and prepared. To date, it has been almost impossible to perform gene expression analysis using very small FFPE tissue sections (diameter: 100 μm) as starting material. In this study, we reported gene expression analysis using very small FFPE tissue sections (diameter: 100 μm) as starting material, which has not been conducted with significant success to date.

Lastly, the application of this method to clinical specimens showed that it is possible to detect genes even in tissue samples stored at room temperature for 6 years after embedding. Although it is expected that the gene groups identified differ based on the tissue sections used and the microenvironmental conditions, we believe the reliability of the detection results in this report to be considerably high, because the accuracy and sensitivity were evaluated carefully in a model case using mice and applied to clinical specimens. Cancer tissue does not only contain sections that can be clearly distinguished into tumor and non-tumor cells. It is also assumed that gene expression may differ between non-tumor areas adjacent to and distant from the tumor site. Evaluation of spatial gene expression in such microenvironments may help improve the understanding of each characteristic morphological site without the use of valuable clinical specimens at the whole tissue section level, which could help elucidate the mechanisms underlying cancer invasion and drug resistance. We hope that our proposed method will find applications in pathology, drug discovery evaluation, and associated fields of medicine, in which transcriptome analysis of tissue preserved for long time periods is often necessary.

## Methods

### Tissue sources

All mice used in this study (ICR, male, age > 2 months, Tokyo Laboratory Animals Science Co. Ltd., Tokyo, Japan) were treated according to protocols approved by the Committee for Animal Experimentation of the School of Science and Engineering at Waseda University (No.2017-A056, 2018-A067) and the law enforced (No. 105) and notification provided (No. 6) by the Japanese government.

Clinical samples were acquired from surgically resected tumors at the Department of Human Pathology, Juntendo University School of Medicine. Immediately after acquisition, the tissues were fixed in 10% neutral-buffered formalin for 24 hours at room temperature, embedded in paraffin after routine processing. Tumor tissue specimens were collected and analyzed under a protocol approved by the institutional review boards of Juntendo University (No.2018090) and Waseda University (No.2017-G001). Informed consent was obtained from the participants or their legal representatives. The study conformed with the Helsinki declaration (1964) and its amendments or comparable ethical standards.

### Preparation of FF tissue sections

Livers resected from the mice were immediately washed with cooled phosphate-buffered saline (PBS, pH 7.4, Thermo Fisher Scientific), immersed in SCEM solution (Leica), and rapidly frozen in liquid nitrogen. The embedded frozen tissues were stored at -80 °C until use in experiments. For total RNA extraction (from bulk samples), thin sections (thickness: 20 μm) were used. For micro-region sampling, 20 μm-thick sections were transferred to cryofilm (SECTION-LAB, Hiroshima, Japan) and immersed and fixed in 99.5% ethanol for 10 s.

### Preparation of FFPE tissue sections

Liver tissues resected from the mice were immediately washed with cooled PBS, immersed in 4% paraformaldehyde with PBS (Nacalai Tesque, Inc., Kyoto, Japan), and fixed at room temperature for 24 h. They were then immersed in 0.1 mol/L-PBS (pH 7.4, Nacalai Tesque, Inc.). After fixation, the tissues were dehydrated by soaking in 70%, 80%, 90%, and 95% ethanol solution for at least 30 min, followed by soaking in 99.5% ethanol solution for at least 30 min; the step was repeated four times, and the solution was changed each time. Hemo-Clear solution (FALMA, Tokyo, Japan) was used as an alternative to xylene for paraffin embedding. Paraplast (Merck, Darmstadt, Germany), a low-melting-point paraffin, was used for embedding.

### Collection of micro-tissue using the semi-automated micro-tissue punching system

The micro-tissue collection device was fabricated by Frontier Biosystems (Tokyo, Japan). The sampling unit is designed such that it can be easily attached to a commercially available microscope. The device consists of a collection unit containing a tissue collection needle, a pumping unit for ejecting the punched tissue, and an injector unit that controls fluid pumping. The punched micro-tissues are ejected into an eight-micro tube strip. The site to be sampled is selected by observing the tissue under a microscope. Multiple locations can be automatically sampled by specifying the sampling site and the tube to be dispensed in the original system installed in the device.

For micro-tissue sampling, a hollow needle made of stainless steel, purchased from CASTEC (Kanagawa, Japan), was used. The tip of the sampling needle was knife-edged for smooth cutting of tissue sections. This sampling needle was connected to the injector via a polytetrafluoroethylene tube (Nichias, Tokyo, Japan). The tube was filled with a 99.5% ethanol solution. The internal parts of the collection needles were pre-washed with 70% ethanol, RNaseZap (Thermo Fisher Scientific), and 99.5% ethanol. A plastic Petri dish with a diameter of 35 mm was placed on the sample table. A 0.05 mm-thick silicone sheet (Kenneth, Osaka, Japan) was placed in the center of the Petri dish, and a cryofilm on which thin tissue sections were previously transferred was placed, with the surface of the tissue facing upward. In micro-tissue sampling, the tissue, the cryofilm on which the tissue was transferred, and the silicon sheet underlying it are captured together. The dispensed solution contained 99.5% ethanol. This was to inhibit RNA degradation. Tissue sections for micro-tissue sampling were stained with HE and then used. FF thin sections were stained after rapid fixation with 99.5% ethanol, whereas FFPE thin sections were stained after deparaffinization.

### Total RNA extraction from a tissue section

Extraction of total RNA from FF or FFPE thin sections was performed using the RNeasy Mini Kit (QIAGEN, Hilden, Germany) or the RNeasy FFPE Kit (QIAGEN), respectively, according to the manufacturer’s instructions. RINs of the extracted RNA were determined using TapeStation 4200 (Agilent, Tokyo, Japan), and concentration was determined using Qubit (Thermo Fisher). The samples were stored at -80 °C until use.

### cDNA library preparation

Total RNA or a micro-tissue was used as a template for the cDNA library. Micro-tissue sections were dispensed with 99.5% ethanol during sampling and stored intact at -80 °C. A vacuum evaporator was used for cDNA library preparation to completely remove ethanol, followed by the subsequent reaction.

For cDNA library preparation, as the first step, the cell membranes were lysed with proteinase K, and the poly(A) RNA was purified using oligo(dT) magnetic beads (referred to as PPP), as previously described. Conducting the PPP process can reduce the carriers of non-coding RNA and increase the ratio of PCGs against the sequence-reads. For FFPE tissues, conducting the process at 85 °C with 15 min of incubation followed by proteinase K treatment ensured the dissociation of the methylation crosslink. When purified total RNA was used as the starting material, 1 μL of total RNA diluted to an appropriate concentration was used. When micro-tissue sections were used as the starting material, ethanol was completely removed, as described above, and used for subsequent reactions. For each sample, a proteinase K reaction solution containing 5 μL of PKD buffer (QIAGEN) and 0.31 μL of ProK (QIAGEN) was added and incubated for 1 h at 56 °C. The solution was then mixed thoroughly with vortexing, spun down, and incubated at 85 °C for 15 min. Oligo(dT) magnetic beads (Dynabeads Oligo (dT)25, 61002, Thermo Fisher) used for mRNA purification were pre-washed three times with equal volumes of 1× hybridization buffer (2× SSPE, 0.05% Tween20, and 0.0025% RNase inhibitor) and then suspended in half the volume of 2× hybridization buffer, in accordance with the instructions provided in the manual. The bead mixture solution was heated using a thermal cycler at 56 °C for 1 min, and then slowly cooled to 25 °C and allowed to stand for 10 min. The reaction mixture was then placed on a magnetic stand, and the supernatant was removed. The magnetic beads were washed twice with 100 μL of ice-cold 1× hybridization buffer and then washed with 100 μL of ice-cold 1× PBS supplemented with 0.0025% RNase inhibitor (Thermo Fisher), following which the wash solution was removed using a magnetic stand. The magnetic beads in which RNA was trapped were resuspended in 2.8 μL of RNase-free water. The RNA bound to the complementary strands of poly(T) on magnetic beads was then denatured by heating at 80 °C for 2 min on a heat block. Finally, 2.5 μL of the supernatant containing purified mRNA was recovered using a magnetic stand and used as material for cDNA library preparation. The total lysed and purified samples were directly processed according to the Smart-seq2 flow. The total mRNA produced during the PPP step was used for cDNA library preparation. The library preparation process was conducted according to the Smart-seq2 flow. The cDNA libraries were produced by adding ERCC at the same concentration (1:120,000 diluted ERCC spike-in mix) to all samples as an internal standard.

### Statical analysis

Statistical analyses were performed using R studio (ver 1.3.959) under R version 4.1.2 environment. The means of two independent groups were compared using Welch’s t-test. The p-values were calculated and visualized using the R package “ggpubr”^33^ based on “ggplot2”^34^.

### Sequencing and data analysis

Amplified cDNA (0.25 ng) was used for preparing the sequencing library with the Nextera XT DNA Library Prep Kit (Illumina, San Francisco, CA, USA). Paired-end sequencing was performed on the MiSeq platform, with 75 bases each for read 1 (R1) and read 2 (R2). We trimmed the adapter sequences in all the sequence reads using Flexbar (ver. 3.5.0). The trimmed sequence reads were aligned to the Ensembl mouse reference genome (GRCm38 ver.92) for mouse liver tissue samples, including the ERCC sequences, using Hisat2 (ver. 2.1.0), with the default parameters. The gene expression levels, expressed as TPM, were calculated using Stringtie (ver. 2.1.7), with a transcriptome reference obtained from Ensembl. For comparison of the expression levels, the counts of mapped reads were normalized by the median of ratios in DESeq2^35^.

## Acknowledgements

We thank Naoko Suzuki and Kiyofumi Takahashi for providing technical support. This research was supported by Platform Project for Supporting Drug Discovery and Life Science Research (Basis for Supporting Innovative Drug Discovery and Life Science Research (BINDS)) from AMED under Grant Number JP21am0101104. The super-computing resource was provided by the Human Genome Center (University of Tokyo).

## Author contributions

H.M., K.A. and H.T. conceived and designed the experiments. H.M., K.A., M.Y., M.H. and H.K. conducted the experiments and collected the data. H.M., K.A., M. Y., R.W. and K.I. performed computing analysis of the results. K.T., K.S., and T. H. collected and analyzed pathological data. H.M., S.A.Z., M.H. and H.T. wrote the manuscript.

## Data availability statement

RNA-seq data were deposited in the Sequence Read Archive (https://www.ncbi.nlm.nih.gov/sra) under the accession number PRJNA815867.

