## Supplementary Figure 1-3 for "Reproducible and sensitive micro-tissue RNA-sequencing from formalin-fixed paraffin-embedded tissue for spatial gene expression analysis"

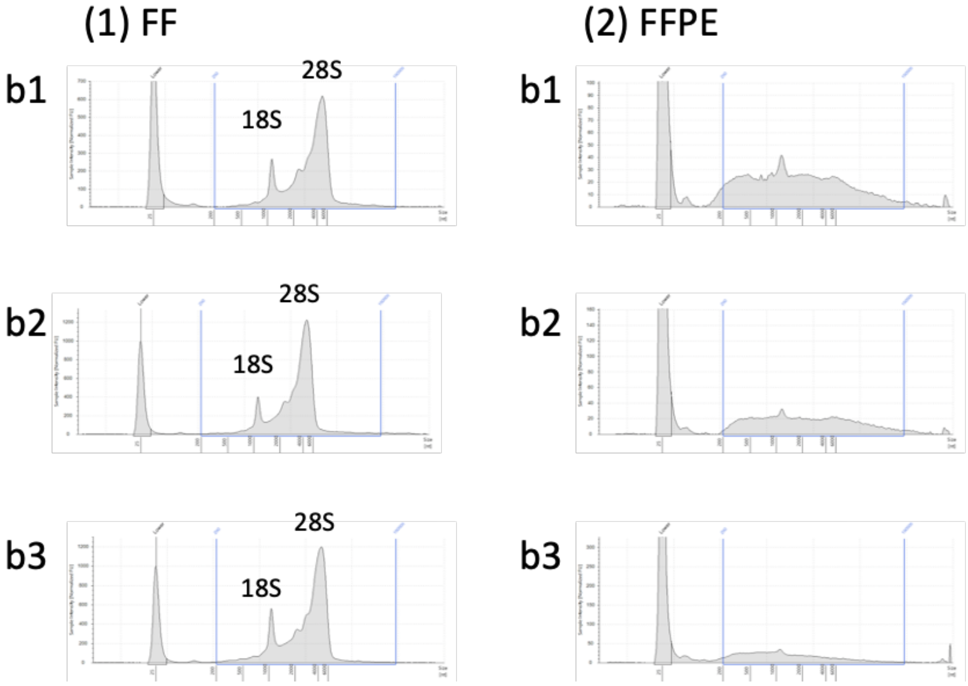


**Supplementary Figure 1. Electropherogram of total RNA**

The electrophoretic patterns of purified total RNA were obtained using a commercially available kit (RNeasy Mini Kit (QIAGEN, Hilden, Germany) or RNeasy FFPE Kit (QIAGEN)). Peaks of ribosomal RNA (18S and 28S) could be detected in total RNA extracted from FF tissues, but not in that from FFPE tissues. Compared with that in FF tissues, total RNA from FFPE tissue was spread uniformly over a wide range of base lengths, from long to short, indicating that the latter was degraded.

FF, fresh-frozen; FFPE, formalin-fixed paraffin-embedded.

Supplementary Figure 2

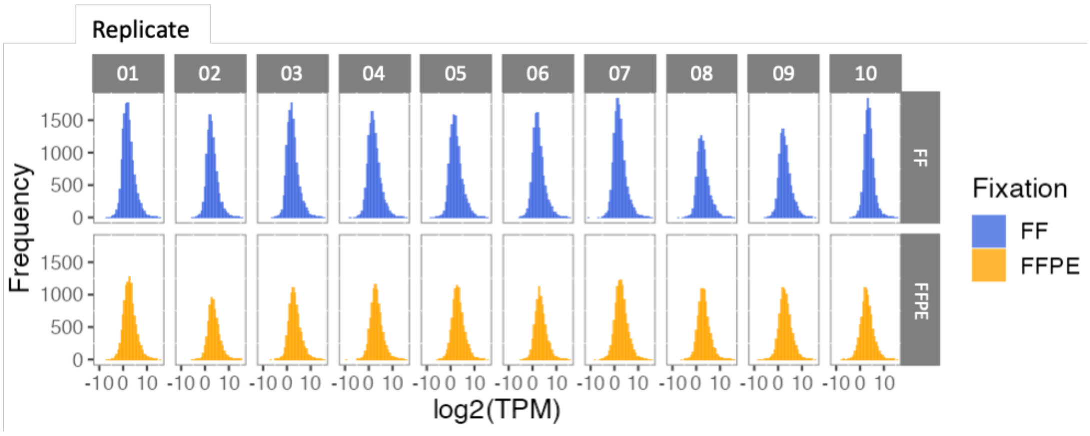


**Supplementary Figure 2. Correlation of gene expression patterns**

The frequency patterns of gene expression in the PuTi-spots (N = 10) obtained from FF and FFPE tissues were highly similar in form, except that the frequency was higher in FF tissues than in FFPE tissues. The frequency of expression is shown on the vertical axis, and the expression (log2(TPM)) on the horizontal axis.

PuTi, micro-punched tissue spot; FF, fresh-frozen; FFPE, formalin-fixed paraffin-embedded.

### Supplementary Figure 3_1

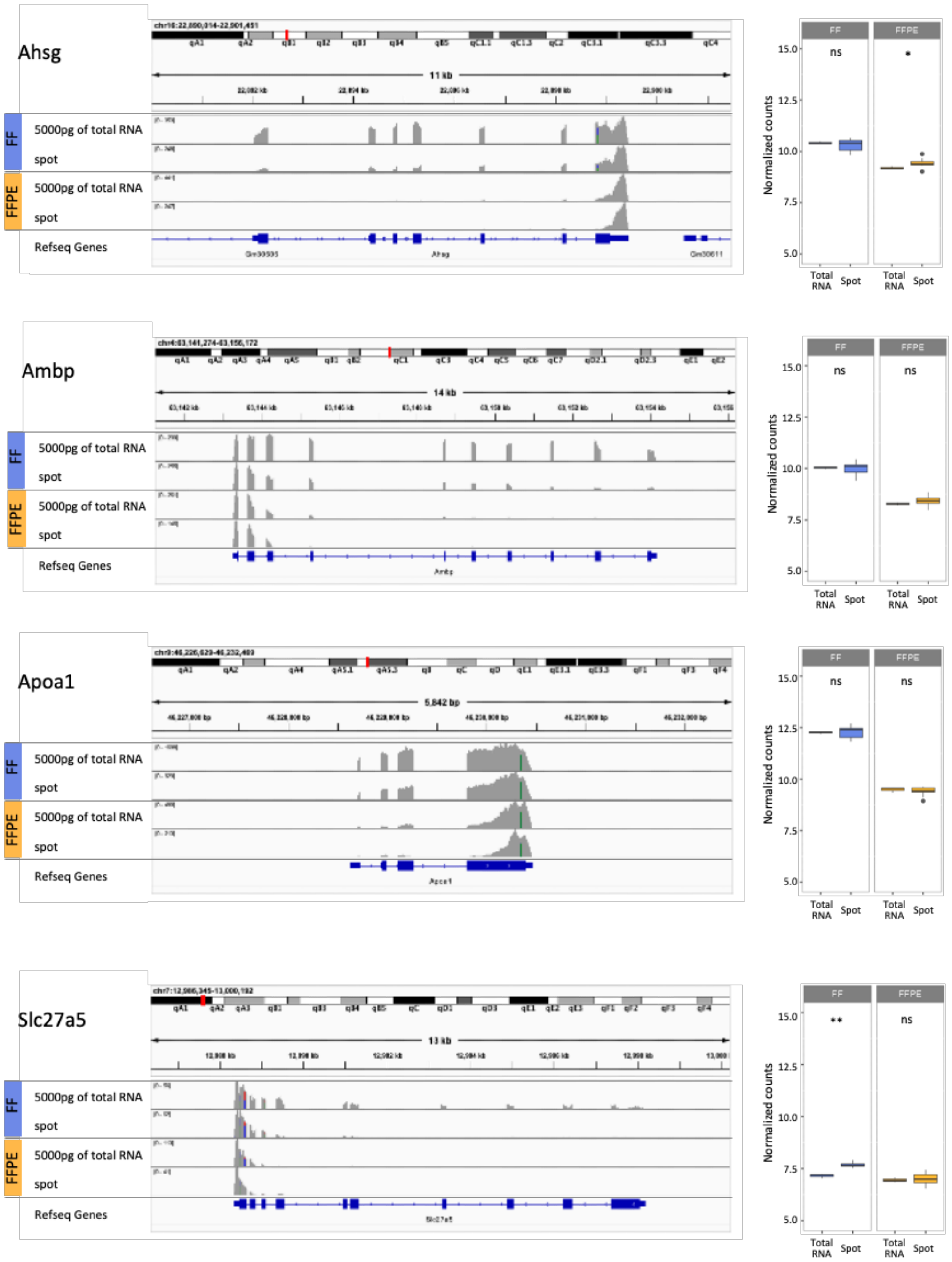

### Supplementary Figure 3_2

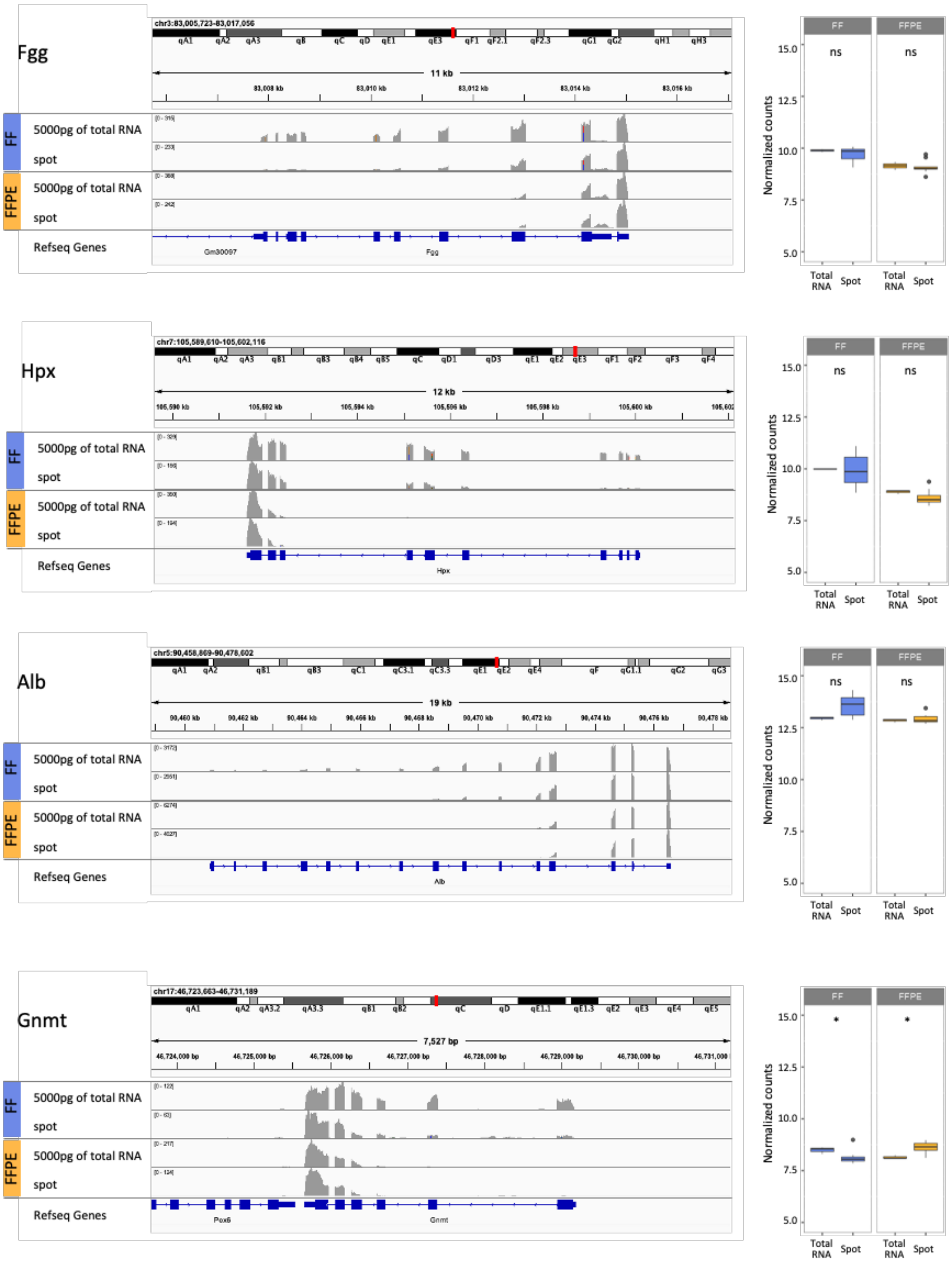

### Supplementary Figure 3_3

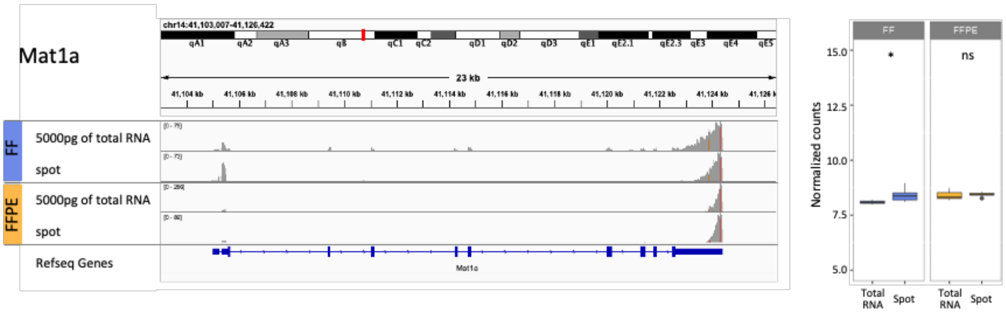


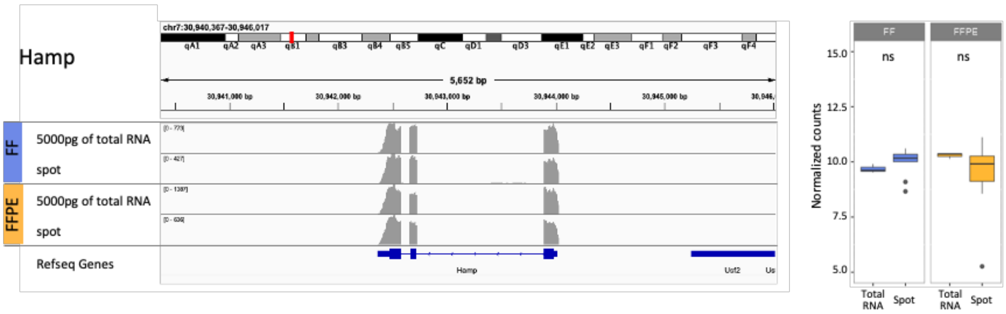

**Supplementary Figure 3. Gene expression comparison between bulk tissue and the tissue spots**

Alignment of reads to ten liver-specific genes (left), and comparison of normalized read counts between purified total RNA and PuTi-spots derived from FF or FFPE tissue (right box-plot). Read coverage was visualized using Integrative Genomics Viewer. The bottom of the column shows visualized exons (bold line and boxes) and introns (line). Sequencing reads derived from purified total RNA are more well-characterized for alignment, depending on the tissue preservation method. A greater number of reads were aligned at the 3'-end of each gene, and fewer reads were mapped toward the 5'-end. In both FF-derived samples (purified total RNA and PuTi-spots), the aligned region of reads was longer and extended toward the 5'-terminus. The difference in normalized read counts between FF and FFPE tissues could be attributed to the length of the alignment from the 3'-end.

PuTi, micro-punched tissue spot; FF, fresh-frozen; FFPE, formalin-fixed paraffin-embedded.

#### Supplementary Figure 4

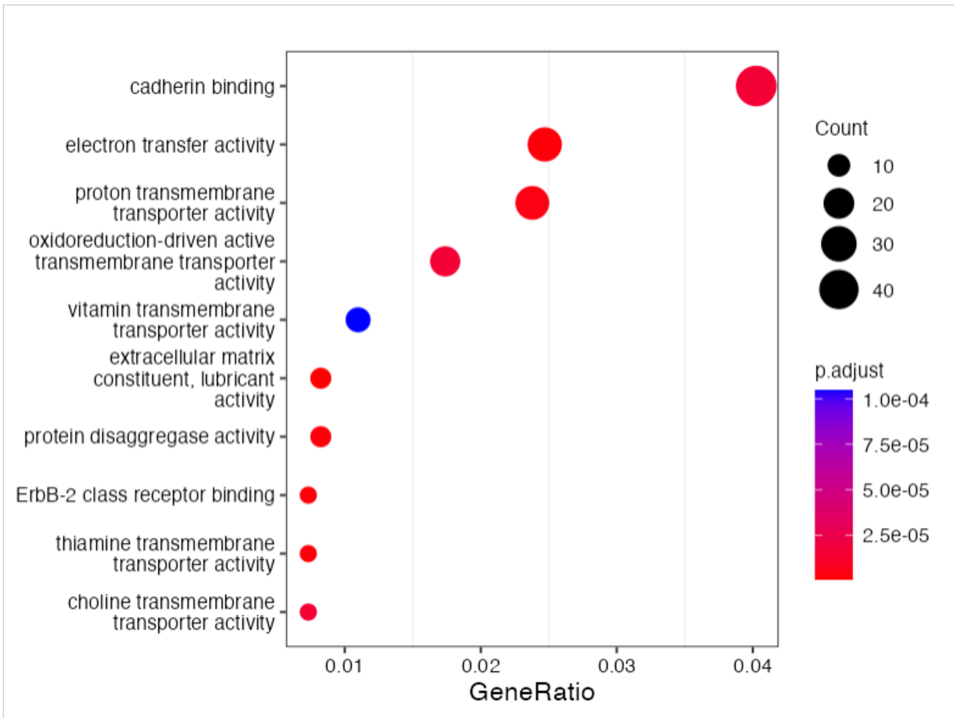


**Supplementary Figure 4.** Gene ontology enrichment analysis of upregulated genes in PuTi-spots collected from the tumor area. Molecular function-linked cadherin binding was enriched in tumor PuTi-spots (*p-adjust* = $1.51 \times 10^{-5}$), and the *S100P* and *TAGLN2* genes were included (Supplementary Table 1).

PuTi-spot, micro-punched tissue spot.
